# Stability and Performance of Linear Combination Tests of Gene Set Enrichment for Multiple Covariance Estimators in Unbalanced Studies

**DOI:** 10.1101/2025.01.23.634558

**Authors:** Sara Khademioureh, Payam Amini, Erfan Ghasemi, Paul Calistrate-Petre, Saumyadipta Pyne, Irina Dinu

## Abstract

Gene set analysis (GSA) is essential for understanding coordinated gene expression changes within biological pathways, especially in high-dimensional data generated by platforms such as RNA-seq and microarrays. This study focuses on the linear combination test (LCT), a GSA method that combines multiple genelevel statistics into a powerful test statistic to assess the association between a gene set and outcomes of interest in a given set of samples.

We evaluated the performance and stability of LCT using different covariance matrix estimators, including ridge, graphical lasso, and adaptive lasso, which are known for their effectiveness in high-dimensional data analysis. In addition, we assessed the robustness of LCT in the face of unbalanced study designs, which are typical in biomedical research due to limited sample availability and the high cost of data generation.

We conducted a simulation study and applied LCT to publicly available gene expression datasets comparing patients with systemic lupus erythematosus (SLE) to healthy controls, where the number of controls is significantly lower than the number of cases. Our findings demonstrate that while LCT’s default shrinkage estimator shows limitations in highly correlated and unbalanced designs, ridge estimation provides a more reliable alternative for unbalanced scenarios. Researchers can optimize LCT’s performance by selecting appropriate covariance estimators based on their data structure. These results suggest that LCT is a reliable and powerful tool for GSA in unbalanced studies, identifying SLE-relevant gene sets more effectively than other GSA methods and showing validation against clinical phenotypes, offering valuable insights into the underlying mechanisms of complex diseases such as SLE.

## 1 Introduction

Microarray and RNA-seq technologies have revolutionized our understanding of the gene expressions in an organism under a given condition. Yet, analyzing such expression data for individual genes can overlook the coordinated activity of genes as it occurs within biological pathways. This limitation hinders our ability to fully grasp the underlying biomedical mechanisms such as of complex diseases. Gene set analysis (GSA) has emerged as a powerful tool for analyzing high-dimensional gene expression data, mostly from microarrays and RNA-seq platforms, by addressing this challenge. It analyzes coordinated changes in functionally related genes within pre-defined gene sets, thus providing a more comprehensive understanding of biomedical phenomena as compared to analyses that focus on individual genes.

Dinu et al. (2007) made a significant contribution to the field of GSA by introducing the significance analysis of microarrays for gene sets (SAM-GS). This method addressed limitations in the platform gene set enrichment analysis (GSEA) (Subramanian et al., 2005; Mootha et al., 2003), a widely used GSA tool for identifying gene sets statistically significantly associated with a categorical phenotype. Building on the above foundation, Wang et al. (2011) introduced the linear combination test (LCT). LCT offered a novel approach by combining multiple gene-level statistics into a single, powerful test statistic. This innovation allowed LCT to effectively detect subtle yet coordinated changes in gene expression within predefined gene sets, particularly those exhibiting weak individual gene signals.

Since its introduction, LCT has undergone continuous development to enhance its capabilities and address new challenges in GSA, including continuous and multivariate outcomes (Dinu et al., 2013; Wang et al., 2014). Further, Khodayari Moez et al. (2019) extended LCT to the analysis of gene sets in studies where gene expression is measured at multiple time points. LCT was also extended to single-cell transcriptomics (Dinu et al., 2021) as well as other omics (Khodayari et al., 2018). Amini et al. (2023) have recently introduced a novel advancement with the geographically weighted LCT (GWLCT). This version allows researchers to analyze gene sets that are relevant to location-specific distribution of spatial phenotypes in tissues, making it a valuable tool for studies involving spatial heterogeneity, such as intratumor heterogeneity in cancer research.

These advances strengthen LCT’s position as a versatile GSA tool capable of handling diverse data types, including longitudinal and spatial data associated with multivariate or continuous phenotypes. Detecting the association between such phenotypes and high-dimensional gene expression data is challenging for traditional statistical methods due to two key reasons. First, the number of genes (features) is often vastly more than the number of samples (observations), making it challenging to estimate relationships between genes accurately. Second, gene expression measurements, particularly within functionally related gene sets, tend to be highly correlated, further complicating the analysis.

However, the performance of LCT has not yet been rigorously analyzed when it is applied to studies with unbalanced designs, a common occurrence in gene expression analysis, say, for binary outcomes such as disease cases and healthy controls. Unbalanced designs occur when the number of samples in each outcome category differs considerably. This imbalance can introduce bias into the analysis, potentially leading to inaccurate identification of differentially expressed gene sets analysis using LCT. Limited sample availability in rare diseases or specific cell types, technical constraints restricting the number of samples processed simultaneously, and the high cost of high-throughput sequencing for extensive studies all contribute to unbalanced designs in binary phenotype research (Kerr, 2009; Yang et al., 2006). To obtain bio-specimens from healthy individuals as control samples is often difficult. Given the prevalence of unbalanced designs, therefore, evaluation of the robustness of GSA methods, including LCT, under such scenarios is both important and useful.

LCT addresses these issues by incorporating a crucial element of employing the shrinkage covariance matrix estimator of gene expression proposed by Schäfer and Strimmer (2005). This matrix captures the relationships between genes within a gene set, allowing LCT to account for correlated expression patterns and deliver more accurate results in the context of high-dimensional data. This study aims to assess the power of LCT using several well-regarded shrinkage estimators for the covariance matrix: Ridge, graphical lasso, and adaptive lasso. These techniques have gained recognition for their effectiveness in addressing the challenges of high-dimensional data commonly encountered in gene expression analysis.

GSA depends not only on the statistical test of a pre-defined gene set’s enrichment in transcriptomic data but also on the quality and availability of such gene sets. There exist several manually curated catalogs of gene sets that are known to be functionally related or act as mechanisms of disease in various organisms. In this study, we utilized the Kyoto Encyclopedia of Genes and Genomes (KEGG) (Kanehisa et al., 2012). KEGG is a highly regarded resource for gene sets and molecular pathways that was initiated in 1995 (Kanehisa and Goto, 2000). It is a comprehensive database that integrates diverse molecular datasets across different omics to map molecular interactions and reactions. KEGG’s pathway maps, BRITE hierarchies, and KO (KEGG Orthology) system facilitate understanding gene functions and their roles in complex networks. Additionally, KEGG includes databases for diseases and drugs, enhancing research into disease mechanisms and drug interactions (Kanehisa et al., 2021). Other gene set catalogs, such as BIOCARTA and Human Phenotype Ontology, were also used.

The present study aims to demonstrate LCT’s ability to maintain reliable and accurate results under unbalanced conditions and different covariance matrix estimators. Towards this, we performed GSA on publicly available, published gene expression datasets generated using different high-throughput platforms for comparing patients with the autoimmune disease systemic lupus erythematosus (i.e., SLE, or simply, lupus) and corresponding healthy controls. As is common among biomedical studies, the controls are considerably fewer in number than the cases, which makes such studies unbalanced. In the following section, we describe the data and methodology used for LCT. In particular, the details of a simulation study with the choice of different covariance matrix estimators and GSA methods are provided. Validation of LCT results was demonstrated based on clinical phenotypes of SLE. We end with a description of the results of our analysis, followed by a general discussion.

## 2 Methods and Data Description

We begin by introducing LCT for binary phenotype, which uses shrinkage as its default covariance matrix estimator to analyze gene expression data. Next, we present alternative estimators for the covariance matrix, including ridge, graphical lasso, and adaptive lasso, which were selected to replace the shrinkage estimator in LCT based on their demonstrated ability to control Type I error in high-dimensional settings and their theoretical appropriateness for analyzing gene expression data. The implementation of these different estimators allows us to evaluate both the computational performance and statistical robustness of LCT across various scenarios of transcriptomic data analysis.

### 2.1 LCT

LCT tests if any linear combination of expressions of genes in a gene set is associated with values of a given phenotype. It identifies the most significant linear combination of genes by maximizing the test statistic derived from the covariance matrix of gene expressions.

Consider gene expression data consisting of a total of *m* genes and *n* subjects (or samples), where the samples are divided into *n*_1_ cases and *n*_2_ controls (*n* = *n*_1_ + *n*_2_) representing two distinct groups. Let *X* = *X*_*ij*_ be the gene expression matrix where *X*_*ij*_ represents the expression of gene *i* for subject *j*. A predefined gene set comprises *k* genes with expressions denoted as {*x*_1_, *x*_2_, …, *x*_*k*_} where *x*_*i*_, *i* = 1, 2, …, *k* refers to the observed gene expression values for the *i*-th gene across all samples. The null hypothesis to be tested is *H*_0_: No linear combination of {*x*_1_, *x*_2_, …, *x*_*k*_} is associated with the phenotype. Let *Z*(*β*) = *β*_1_*X*_1_+…+*β*_*k*_*X*_*k*_ be a linear combination of the genes.

For a given vector *β* = (*β*_1_, *β*_2_, …, *β*_*k*_)^*T*^ of combination coefficients, we test whether the combination *Z*(*β*) is associated with the phenotype using the following univariate model:

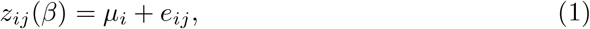

where *µ*_1_ and *µ*_2_ are the mean gene expressions for cases and controls, respectively, and *e*_*ij*_ ~ *N* (0, *σ*^2^), with *j* denoting 1, …, *n*_*i*_ of group *i* = 1, 2.

The test statistic for *Z*(*β*) is based on the two-sample *t*-test, *T* (*β*), which follows a t-distribution, or equivalently *T* ^2^(*β*) follows an F-distribution. To test *H*_0_, we consider the most significant linear combination, *β*^*∗*^, among all possible combinations is found by maximizing *T* ^2^(*β*):

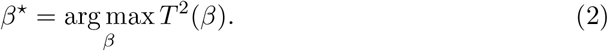

As the square of the two-sample *t*-test, we have:

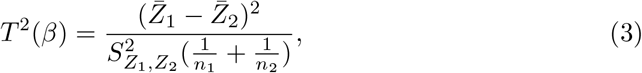

where 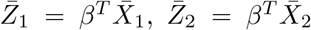 are the sample means of *Z*(*β*) for cases and controls respectively, 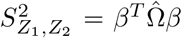 is the pooled sample variance, 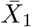 and 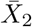 are *k*-sample averages of gene expressions within cases, and controls respectively, and 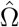 is the estimated pooled covariance matrix over the two phenotype groups.

When the size of the gene set *k* is larger than the sample size *n*, the covariance matrix Ω is singular. To address this, LCT employs a shrinkage covariance matrix proposed by Schäfer and Strimmer (2005), replacing the singular covariance matrix 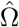, with a shrinkage covariance matrix 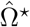to handle high-dimensional data.Let 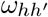 denotes the (*h, h*^*′*^)-th entry of 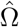, which is defined as:

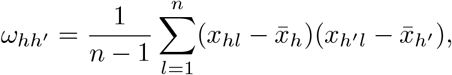

where *x*_*hl*_ is the expression of gene *h* in sample *l*, and 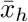 is the mean expression of gene *h* across all samples. Then the shrinkage covariance matrix 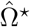 is given by which is a pooled correlation coefficient for the *hh*^*′*^-th gene:

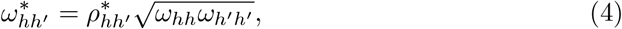

with shrinkage coefficients 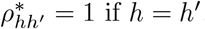 and 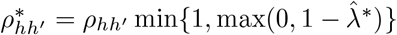 if *h* ≠ *h*^*′*^, where 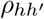 is the pooled sample correlation between the *h*-th and *h*^*′*^-th genes. The optimal shrinkage intensity is estimated as:

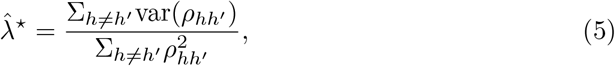

To enhance computational efficiency, we employ a set of normalized orthogonal bases in place of the original observation vectors. This involves conducting an eigenvalue decomposition of the shrinkage covariance matrix of 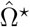:

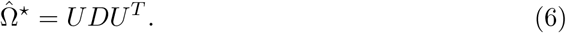

We then obtain a group of orthogonal basis vectors 6. The square of the two-sample test statistic can be rewritten as: 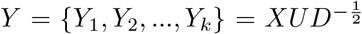

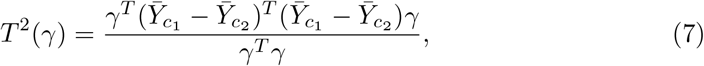

where 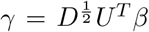 and 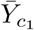 and 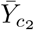 are the *k*-sample means of *Y* for cases and controls respectively. The coefficients of the most significant combination are given by:

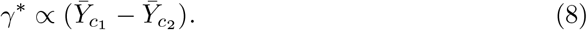

The LCT statistic is proportional to the *L*_2_-norm of the mean-vector difference between the two groups after the orthogonal transformation:

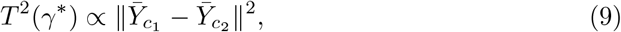

The statistical significance against the null hypothesis is evaluated using a permutation test (i.e., by permuting phenotypic group labels). This approach is computationally advantageous, as the eigenvalue decomposition of 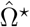 is performed only once, rather than for each permutation. The shrinkage method adjusts the covariance matrix by shrinking the off-diagonal elements toward zero, controlled by the shrinkage intensity *λ*^*∗*^. This also helps manage multicollinearity and ensures that the covariance matrix is invertible.

The shrinkage approach is the default method in LCT for estimating the covariance matrix, and we used the ‘cov.shrink’ function from the ‘corpcor’ R package (Schafer et al., 2021). The estimation was performed with variance shrinkage intensity set to zero where the empirical variances are recovered, and optimal correlation shrinkage to compute the covariance matrix estimate 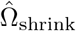 for use in the LCT framework, as described in (Opgen-Rhein and Strimmer, 2007). This approach applies optimal shrinkage to correlations, allowing the method to automatically determine the optimal shrinkage intensity for the correlation estimates while maintaining the original variance structure of the data.

### 2.2 Alternative Estimators for the Covariance Matrix

In this study, we evaluate three alternative covariance matrix estimators within the LCT framework: ridge, graphical lasso, and adaptive lasso, comparing them with the default shrinkage estimator for Ω^*⋆*^. When analyzing high-dimensional gene expression data where the number of genes often exceeds the sample size, accurate estimation of the covariance matrix becomes challenging due to potential singularity and instability issues. Based on our preliminary experiments, these alternative estimators demonstrated strong Type I error control and acceptable statistical power while being theoretically appropriate for high-dimensional data analysis. Through simulation studies with diverse scenarios and real-world data applications, we evaluated the performance and robustness of these estimation methods.

#### 2.2.1 Ridge Estimator

For the ridge estimation approach, we used the ‘ridgeP’ function from the ‘rags2ridges package R (Peeters et al., 2022) to estimate the covariance matrix, which employs a proper L2-penalty (Van Wieringen and Peeters, 2016). As *λ* represents the value of the penalty parameter in this function, the estimation was performed using *λ* = 1 to calculate the estimate of the covariance matrix 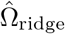 for use in the LCT framework. This choice of *λ* was optimized through our simulation studies, although larger values might be needed depending on the specific data structure and dimensionality and very small values would provide insufficient regularization and potentially lead to poorly conditioned estimates in high-dimensional settings.

#### 2.2.2 Graphical Lasso Estimator

For the graphical lasso approach, we used the ‘glasso’ function from the ‘glasso’ R package (Friedman et al., 2008, 2019) to estimate the covariance matrix, which employs an L1 penalty to induce sparsity. As *ρ* represents the value of the penalty parameter in this function, the estimation was performed using *ρ* = 0.4 to calculate the estimate of the covariance matrix 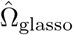 for use in the LCT framework. This choice of *ρ* was optimized through our simulation studies, as very small values would significantly increase the computational time due to too many spurious connections, while larger values might miss important gene relationships.

#### 2.2.3 Adaptive Lasso Estimator

For the adaptive lasso approach, we used the ‘adaptiveLassoEst’ function from the ‘cvCovEst’ R package (Rothman et al., 2009; Zou, 2006) to estimate the covariance matrix, which employs an adaptive L1 penalty. Estimation was performed using a penalty parameter *λ* = 0.5 and an adaptive weight exponent *n* = 0.8 to calculate the estimate of the covariance matrix 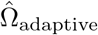 for use in the LCT framework. The choice of *λ* provides a balance between sparsity and bias, helping to maintain stability in high-dimensional settings while avoiding overshrinkage of important coefficients. The adaptive weight exponent *n* = 0.8 was chosen as it allows sufficient differentiation between large and small coefficients, helping reduce bias for large coefficients while maintaining sparsity.

### 2.3 Data Description

In this study, we analyzed three distinct gene expression datasets from the Gene Expression Omnibus (GEO) database to evaluate LCT’s performance in two key aspects: its robustness with highly unbalanced designs and the comparative effectiveness of different covariance matrix estimators for handling matrix singularity. For the first objective of assessing LCT’s performance with notably unbalanced designs, we examined two datasets: GSE99967 (Wither et al., 2018) and GSE72509 (Hung et al., 2015). GSE99967 consists of gene expression microarray data measuring 24,729 genes across 59 samples, with an unbalanced design of 42 SLE cases who met the revised 1997 American College of Rheumatology classification criteria for SLE were included in the analysis and 17 healthy controls (71% cases) based on their whole blood samples. GSE72509 provided RNA sequencing data for 37,105 genes across 117 samples, featuring an even more pronounced imbalance with 99 SLE cases and 18 healthy controls (85% cases) based on their whole blood samples. Both datasets were analyzed using the KEGG Legacy catalog from the Molecular Signature Database (MSigDB) (Human MSigDB Collections, 2023), comprising 186 gene sets.

For the second objective of comparing different covariance matrix estimators, we analyzed GSE38351 (Smiljanovic et al., 2012), an expression profiling by array dataset comprising gene expression measurements of 13,895 genes across 26 samples. This dataset featured a relatively balanced design with 14 SLE cases and 12 healthy controls (54% cases). We evaluated this dataset using the BIOCARTA pathway catalog from MSigDB containing 217 gene sets. Further, we used it to clinically validate LCT results based on SLE phenotypes from Human Phenotype Ontology (Köhler et al., 2021). The use of diverse data types (both microarray and RNA sequencing), different pathway catalogs (KEGG and BIOCARTA), and multiple validation methods (computational and clinical) enabled a comprehensive evaluation of LCT’s capabilities across varying experimental conditions.

## 3 Simulation

Following the simulation design outlined in previous LCT studies (Wang et al., 2014), we conducted a comprehensive simulation study to evaluate the robustness of LCT under various conditions. The primary aims of this study were to (a) assess its performance for unbalanced sample sizes between cases and controls and (b) compare LCT’s default shrinkage estimator with three different alternatives: ridge, graphical lasso, and adaptive lasso estimators to evaluate its performance for high-dimensional gene expression data with singular covariance matrices.

In this simulation, we varied the phenotype sample sizes, with one scenario having 20 subjects (total of cases and controls) representing small sample sizes in transcriptomic studies (Lin et al., 2010), and another with 50 subjects representing large sample sizes. We considered both balanced and unbalanced phenotype distributions. For the balanced studies, we used equal numbers of cases and controls: 10 cases and 10 controls for the small sample size (total n=20) and 25 cases and 25 controls for the moderate sample size (total n=50). For the unbalanced studies, we implemented a 1:4 case: control ratio with 4 cases and 16 controls in the small sample size scenario (total n=20) and 10 cases and 40 controls in the moderate sample size scenario (total n=50). In both scenarios, 80% of participants are controls. These ratios were chosen to reflect commonly encountered imbalances in real biomedical studies. We conducted preliminary analyses with less unbalanced ratios and found similar patterns of results. To reflect varying degrees of gene correlation, we used two correlation coefficients (*ρ*), set at 0.9 to represent a high correlation and 0.1 to represent a low correlation between genes. To evaluate the performance of LCT under varying gene set sizes, we used three different gene set sizes of 50, 100, and 200 genes. Each scenario was replicated 1000 times to obtain reliable statistical estimates. The p-values were computed by testing with 1000 permutations.

LCT is generally applied to normalized gene expression data; therefore, gene expression for 10,000 genes was simulated separately and independently from multiple correlated normal distributions for phenotypic groups i=1,2, where i=1,2 denotes cases and controls, respectively. The mean difference, *γ*, in gene expression levels between cases and controls ranged from 0 to 2 in increments of 0.1. The simulation and application were implemented using RStudio (RStudio team, 2023).

LCT performance assessment involved the calculation of the power and Type I error rates based on 1000 replications for each scenario using a nominal *α* level of 0.05. Table 1 and Figures 1-3 illustrate empirical Type I errors and the power of the LCT in all simulation settings. This analysis revealed that unbalanced designs significantly impact the LCT performance, with this limitation becoming more pronounced as gene set size increased. Using shrinkage estimator in LCT showed inflation of Type I error rates (0.072-0.098) in unbalanced designs with high correlation (*ρ* = 0.9)scenarios, while LCT with ridge estimation demonstrated more stable Type I error control, notably showing better control in unbalanced designs (0.028-0.052) compared to balanced designs (0.059-0.071), but with limited power gain. The LCT using graphical lasso estimator became overly conservative in unbalanced scenarios (Type I error rates as low as 0.007-0.012 for sample size 50), while LCT with adaptive lasso maintained consistent Type I error control (0.034-0.062) but exhibited limited power for detecting smaller effect sizes. The impact of correlation structure emerged as a crucial factor in LCT performance. In high-correlation settings (*ρ* = 0.9), unbalanced designs showed markedly reduced statistical power compared to balanced designs, particularly at n=20. The LCT’s Type I error inflation with shrinkage estimator was most pronounced under these high-correlation conditions while using graphical lasso became extremely conservative. In contrast, under low-correlation settings (*ρ* = 0.1), balanced and unbalanced scenarios maintained more similar power profiles, and the differences between estimators in LCT became less pronounced. The correlation effects were particularly evident in larger gene sets, where LCT performance with graphical lasso deteriorated more severely in high-correlation settings.

**Table 1.** Type I error rates across all scenarios observed in the simulation study (*α* = 0.05).

| Sample Size | Gene Set Size | Design | $\rho$ | Shrinkage | Ridge | Graphical Lasso | Adaptive Lasso |
| --- | --- | --- | --- | --- | --- | --- | --- |
| 20 | 50 | Balanced | 0.9 | 0.051 | 0.059 | 0.062 | 0.049 |
|  |  |  | 0.1 | 0.046 | 0.058 | 0.063 | 0.055 |
|  |  | Unbalanced | 0.9 | 0.098 | 0.039 | 0.245 | 0.055 |
|  |  |  | 0.1 | 0.051 | 0.049 | 0.050 | 0.052 |
|  | 100 | Balanced | 0.9 | 0.047 | 0.069 | 0.055 | 0.041 |
|  |  |  | 0.1 | 0.051 | 0.059 | 0.053 | 0.055 |
|  |  | Unbalanced | 0.9 | 0.082 | 0.033 | 0.262 | 0.054 |
|  |  |  | 0.1 | 0.064 | 0.041 | 0.054 | 0.037 |
|  | 200 | Balanced | 0.9 | 0.046 | 0.071 | 0.047 | 0.061 |
|  |  |  | 0.1 | 0.047 | 0.059 | 0.051 | 0.059 |
|  |  | Unbalanced | 0.9 | 0.076 | 0.028 | 0.276 | 0.062 |
|  |  |  | 0.1 | 0.065 | 0.052 | 0.055 | 0.039 |
| 50 | 50 | Balanced | 0.9 | 0.043 | 0.047 | 0.063 | 0.034 |
|  |  |  | 0.1 | 0.046 | 0.035 | 0.060 | 0.041 |
|  |  | Unbalanced | 0.9 | 0.079 | 0.012 | 0.255 | 0.037 |
|  |  |  | 0.1 | 0.059 | 0.050 | 0.051 | 0.047 |
|  | 100 | Balanced | 0.9 | 0.039 | 0.053 | 0.053 | 0.055 |
|  |  |  | 0.1 | 0.067 | 0.048 | 0.055 | 0.049 |
|  |  | Unbalanced | 0.9 | 0.072 | 0.009 | 0.270 | 0.052 |
|  |  |  | 0.1 | 0.057 | 0.044 | 0.066 | 0.059 |
|  | 200 | Balanced | 0.9 | 0.041 | 0.057 | 0.054 | 0.057 |
|  |  |  | 0.1 | 0.063 | 0.047 | 0.051 | 0.039 |
|  |  | Unbalanced | 0.9 | 0.089 | 0.007 | 0.280 | 0.043 |
|  |  |  | 0.1 | 0.065 | 0.055 | 0.069 | 0.053 |

**Fig. 1.**
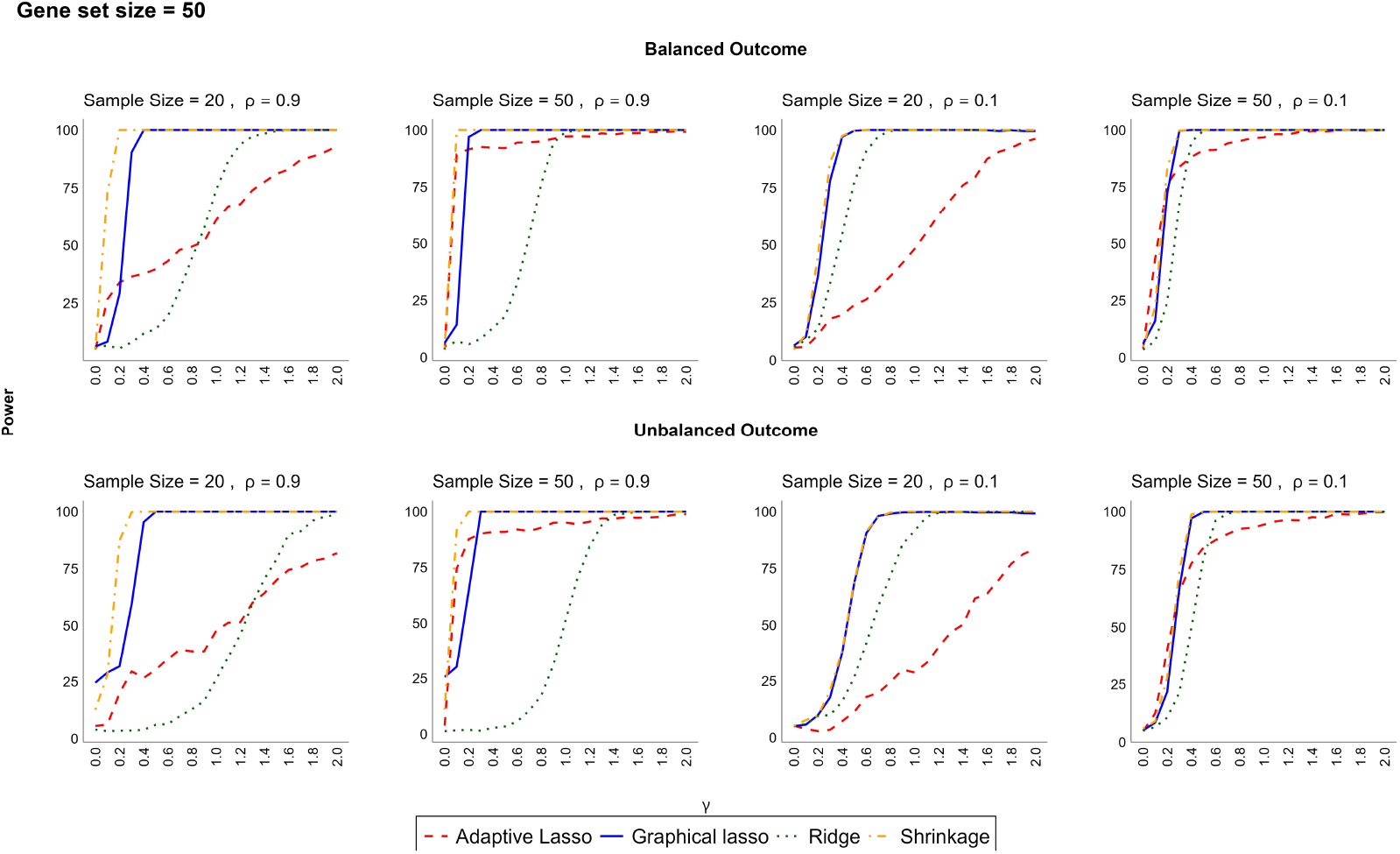
Power (y-axis) comparison of LCT performance for unbalanced and balanced designs using different covariance estimators for gene set size = 50. *γ* (x-axis) is the mean difference in gene expression levels between case and control groups.

**Fig. 2.**
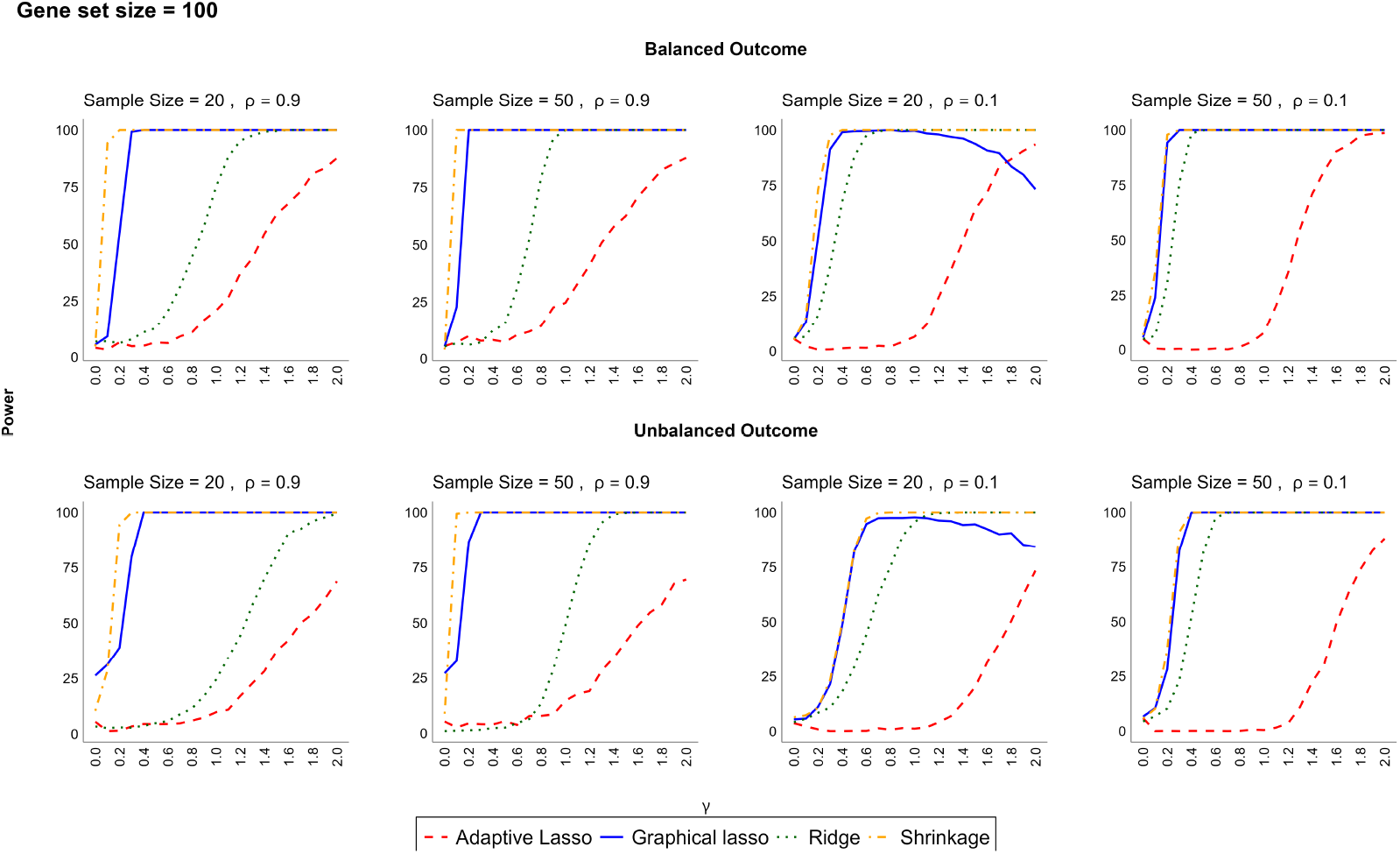
Power (y-axis) comparison of LCT performance for unbalanced and balanced designs using different covariance estimators for gene set size = 100. *γ* (x-axis) is the mean difference in gene expression levels between case and control groups.

**Fig. 3.**
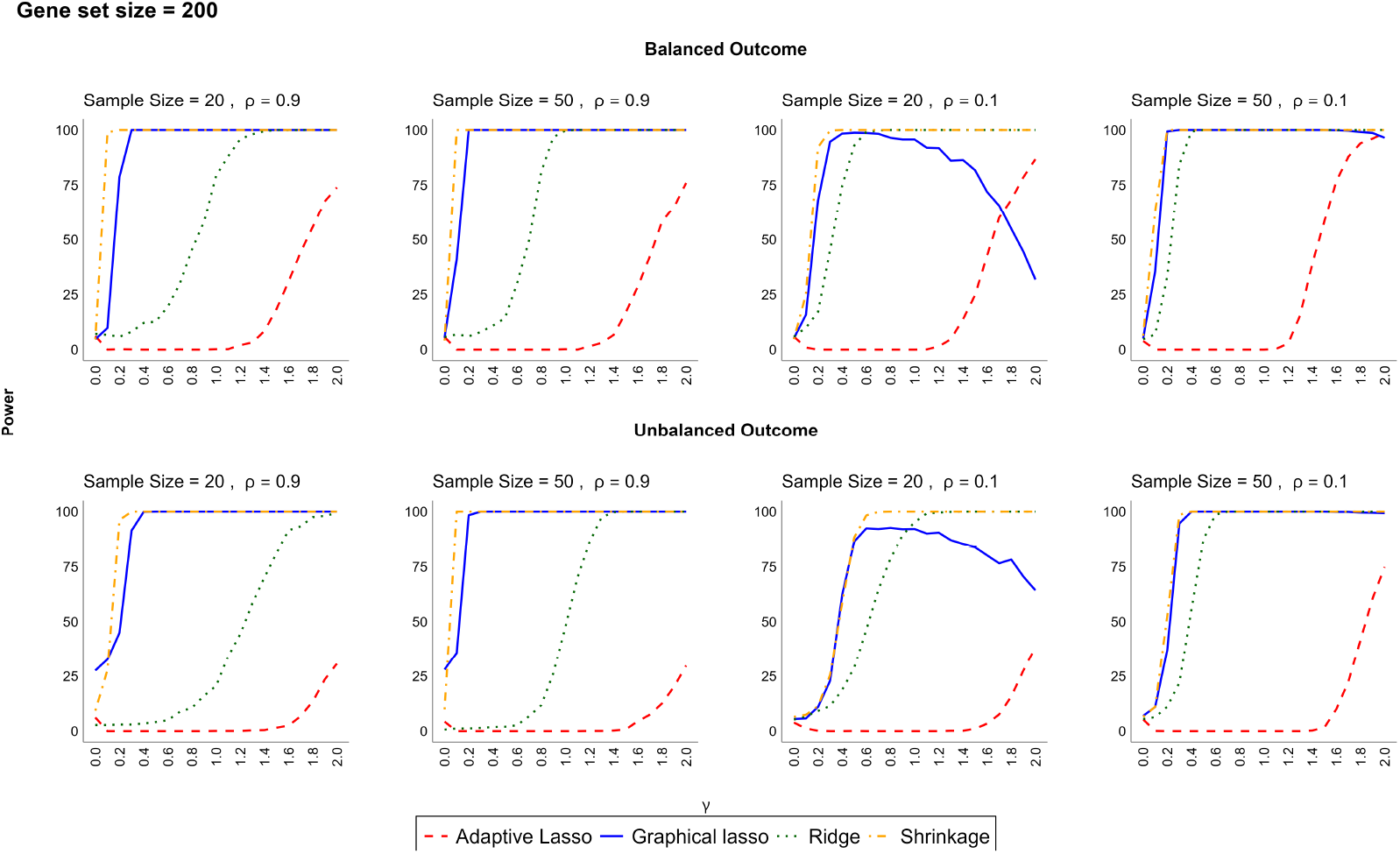
Power (y-axis) comparison of LCT performance for unbalanced and balanced designs using different covariance estimators for gene set size = 200. *γ* (x-axis) is the mean difference in gene expression levels between case and control groups.

Comparing the alternative estimators to the default shrinkage estimator revealed distinct patterns in handling singular covariance matrices, particularly evident as gene set size increased. In balanced designs, where all methods perform well as expected, the shrinkage estimator maintained good power while controlling Type I error, demonstrating its effectiveness for high-dimensional data. However, its performance deteriorated in unbalanced scenarios, particularly with larger gene sets (100, 200), where the covariance matrix becomes more likely to be singular. ridge estimation, while more conservative in power, showed more robust handling of singular matrices across all gene set sizes, making it a potentially safer choice for highly unbalanced studies, especially in the case of high correlation across the set. Graphical lasso showed the most sensitivity to matrix singularity, with its performance declining sharply in larger gene sets, particularly under high correlation. Adaptive lasso demonstrated a middle-ground approach, maintaining Type I error control but with power limitations that became more pronounced with increasing dimensionality.

These limitations and challenges are particularly important for researchers to consider when designing their studies. The results of balanced versus unbalanced designs show substantial power differences, with unbalanced designs (1:4 case: control ratio) showing 20-30% lower power compared to balanced designs when using shrinkage estimator at *ρ* = 0.9. This power reduction was most pronounced for sample size n=20, where the unbalanced design achieved only 40-50% power compared to 70-80% power in balanced designs at the same mean difference in gene expression (*γ* = 1.0). The 1:4 case: control ratio particularly impacted performance with larger gene sets, where power dropped by up to 40% compared to balanced designs.

Based on these comprehensive findings and acknowledging the good performance in balanced scenarios while warning users about unbalanced limitations, we recommend: (1) For balanced designs with high correlation (*ρ* = 0.9), the shrinkage estimator provides optimal performance across all gene set sizes. (2) For unbalanced designs with high correlation, ridge estimation offers the most reliable performance with better Type I error control than in balanced designs, though researchers should be aware of its reduced power for detecting small mean differences in gene expression. (3) For studies with low correlation (*ρ* = 0.1), both shrinkage and ridge estimators perform adequately regardless of balance, with shrinkage showing slight advantages in power. (4) When sample sizes are small (n=20), and designs are unbalanced, ridge estimation is recommended regardless of correlation structure, as it provides the most stable Type I error control. (5) For larger gene sets (100, 200) with unbalanced designs, researchers should consider increasing sample sizes to n=50 or greater to maintain adequate power, particularly when using ridge estimation.

The impact of high dimensionality was particularly evident in unbalanced scenarios, where larger gene sets (100, 200) showed more pronounced performance differences between estimators, reflecting the increased challenge of handling singular covariance matrices with limited sample sizes. These findings emphasize the importance of carefully considering study design, correlation structure, and gene set size when selecting a covariance estimator for LCT analysis, supporting a context-dependent selection strategy. These limitations generally become less pronounced with larger sample sizes (n=50), suggesting that increasing sample size when possible may help mitigate these issues.

## 4 Applications and Results

Following the simulation study, we evaluated LCT’s performance using real high-throughput human gene expression datasets from the GEO repository. We analyzed three SLE datasets that provided different scenarios of sample balance and data types.

For the first objective of assessing LCT’s performance with notably unbalanced designs, we examined two datasets: GSE99967 consists of gene expression microarray data measuring 24,729 genes across 59 samples (42 SLE patients and 17 healthy controls). GSE72509 provided RNA sequencing data for 37,105 genes across 117 samples (99 SLE cases and 18 healthy controls). For comparative analysis, we synthesized balanced versions of our real-world data by randomly selecting cases equal to the number of controls in each study. We repeated this sampling with replacement 15 times to produce 15 synthetic balanced studies from the gene expression data. In each iteration, we conducted GSA and reported the median p-value for each gene set over all 15 iterations as the output of GSA as the balanced design output.

For GSA, we used the KEGG Legacy catalog of 186 genesets from the MSigDB. The biological relevance of each geneset *G* in this catalog to the disease of interest, SLE, was calculated using the well-known Jaccard Index (*JI*). This index provides a baseline measure of similarity between *G* and the curated KEGG gene signature of SLE. To dichotomize, genesets with positive *JI* values are classified as “SLE-relevant,” as opposed to those with *JI* values of zero, which are considered “SLE-irrelevant.”

The application of the LCT on the two transcriptome datasets led to the identification of gene sets associated with SLE, a major autoimmune disease. Moreover, we performed a comparative GSA between LCT and GSEA (a popular GSA method based on the Kolmogorov-Smirnov statistic) using the same datasets and KEGG LEGACY catalog of gene sets for both unbalanced and balanced designs.

The results are shown in Figures 4 (for GSE72509 dataset) and 5 (for GSE99967). The significance score of a gene set *G*, depicted as a point, is transformed by the negative logit of *P*_*G*_ where *P*_*G*_ is the GSA p-value of *G* in the unbalanced scenario. Higher significance scores indicate significant gene sets (p-values *<* 0.05). For the balanced version, the significance score of *G* is computed similarly, but first, GSA is conducted for 15 synthetically balanced scenarios (as described previously), and then their median *P*_*G*_ is used for computing the negative logit of *P*_*G*_. Thus, the smaller the GSA p-value *P*_*G*_ of a gene set *G*, the higher its significance score in the figures. The points in the scatter plots represent gene sets, with x- and y-coordinates showing their relative significance under unbalanced and balanced conditions, respectively. Red points indicate SLE-relevant gene sets, while blue points show SLE-irrelevant ones. The left plots display LCT results, and the right plots show GSEA results.

**Fig. 4.**
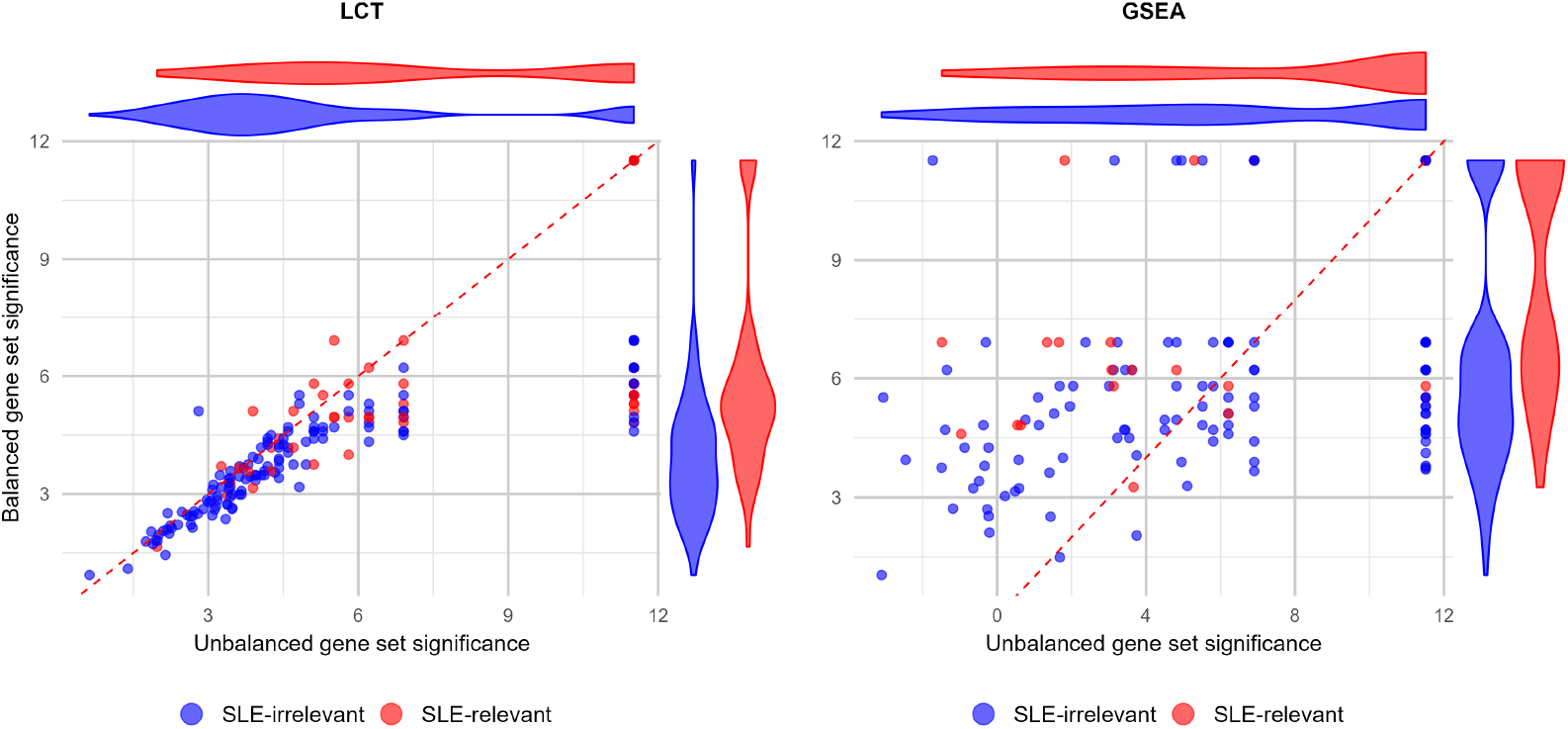
GSA results for the GSE72509 dataset using LCT (left plot) and GSEA (right plot). Each scatterplot displays the negative logit-transformed p-values of SLE-relevant (red) and SLE-irrelevant (blue) gene sets, comparing unbalanced (x-axis) versus balanced (y-axis) scenarios

Interestingly, the markedly higher correlation of the significance scores in the left plots of Figures 4 and 5 underscores that GSA results of LCT are more robust than those of GSEA whether the designs are balanced or not. Further, as described by the marginal violin plots of the two figures for LCT, the SLE-relevant gene sets (red violins) have higher significance values than the SLE-irrelevant ones (blue violins). While this is true for LCT in both studies and under both balanced and unbalanced conditions, no such distinction is noticeable in the violin plots for GSEA in either study.

**Fig. 5.**
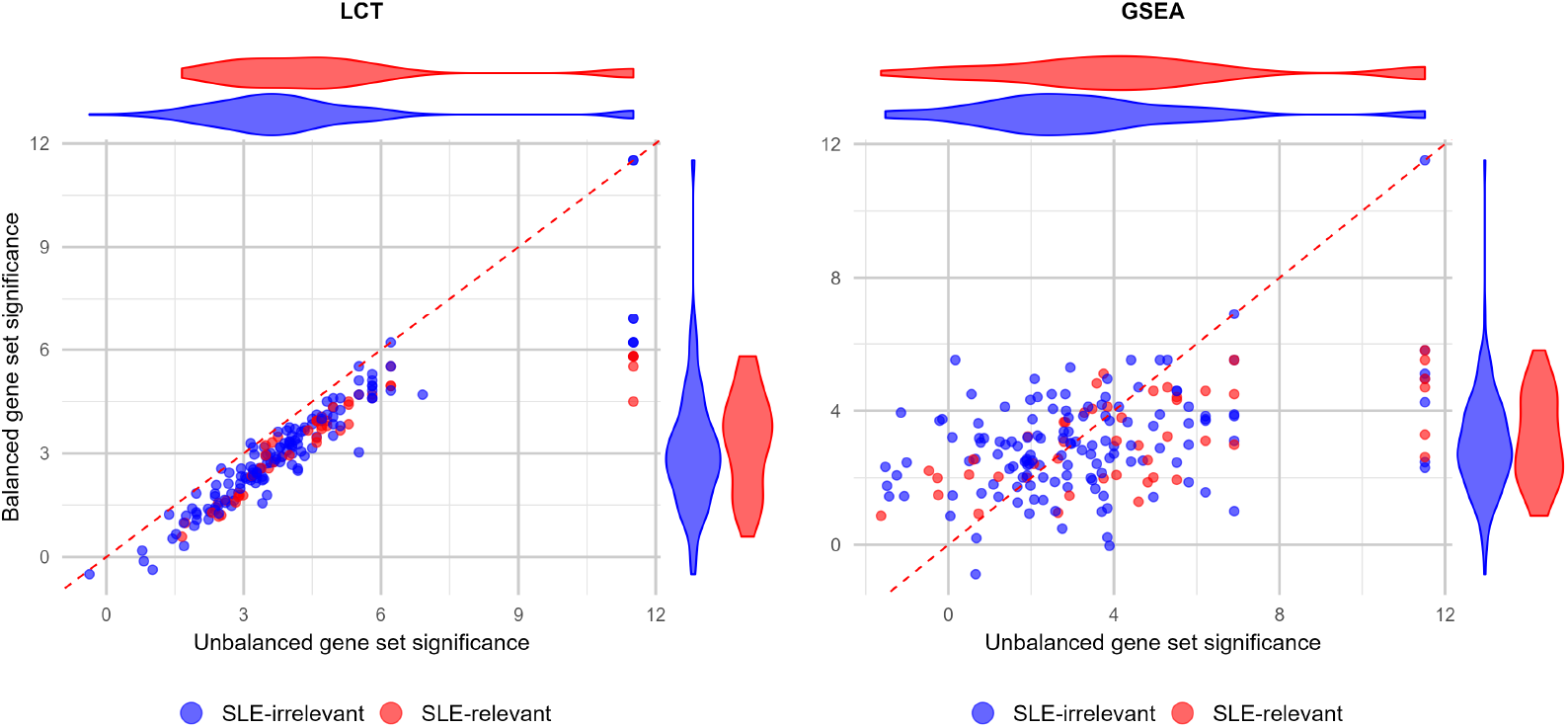
GSA results for the GSE99967 dataset using LCT (left plot) and GSEA (right plot). Each scatterplot displays the negative logit-transformed p-values of SLE-relevant (red) and SLE-irrelevant (blue) gene sets, comparing unbalanced (x-axis) versus balanced (y-axis) scenarios.

However, the notable concentration of significant gene sets, as illustrated by high significance scores (around 12) in the unbalanced scenario, warrants careful interpretation. Given our simulation findings of elevated Type I error rates for LCT in unbalanced designs, these significant gene sets should be interpreted with appropriate caution. While LCT maintains better consistency across different study designs compared to GSEA, users may need to consider more stringent significance thresholds when applying LCT to heavily unbalanced datasets.

Under unbalanced conditions, where the control samples were significantly fewer than the case samples, the LCT methodology robustly identified multiple key pathways. For example, the “KEGG Cytosolic DNA Sensing Pathway” was significantly enriched (p-value *<* 0.001, q-value *<* 0.001), highlighting its potential involvement in the autoimmune responses characteristic of SLE. Additionally, the “KEGG Toll Like Receptor Signaling Pathway”, which is integral to the innate immune system, was significantly enriched (both p-value and q-value *<* 0.001). Such findings illustrate the LCT’s robustness to detect significant associations even when faced with the challenge of outcome imbalance.

For the second objective of comparing different covariance matrix estimators, we analyzed GSE38351, a dataset comprising gene expression measurements of 13,895 genes across 26 samples (14 SLE cases and 12 healthy controls). This relatively balanced dataset was evaluated using the BIOCARTA pathway catalog containing 217 gene sets, allowing us to assess the performance of four different methods (LCT’s default shrinkage estimator against the ridge, graphical lasso, and Adaptive lasso estimators) for identifying significant pathways across various *p*-value thresholds in a real-world context.

The results are shown in Table 2. The shrinkage method demonstrated the highest sensitivity across all significance levels, identifying 249 (85.3%) significant pathways at *p*-value *<* 0.05 and maintaining robust detection with 125 (42.8%) pathways at the most stringent threshold of *p*-value *<* 0.001. The graphical lasso emerged as the second most sensitive method, detecting 225 (77.1%) pathways at *p*-value *<* 0.05 and 95 (32.5%) pathways at *p*-value *<* 0.001, suggesting it provides a moderately conservative approach while maintaining good detection power.

**Table 2.**
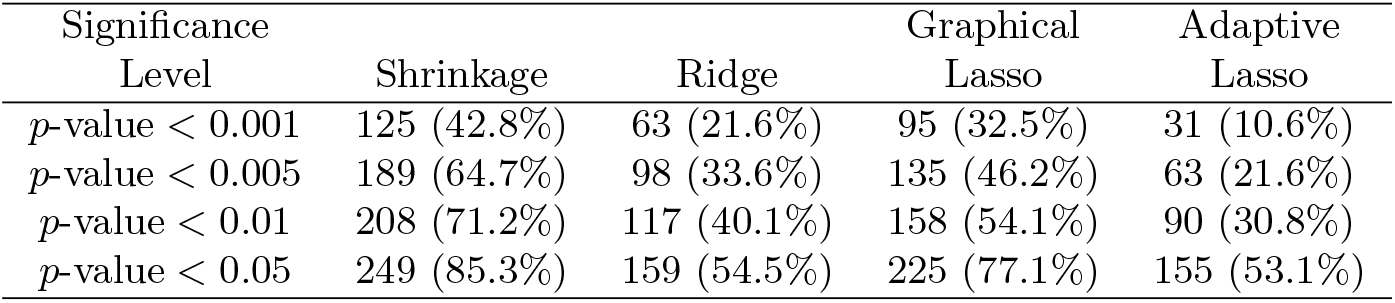
Distribution of results on significant BIOCARTA pathways from four estimators by different p-value thresholds. The results are presented as count (%).

| Significance Level | Shrinkage | Ridge | Graphical Lasso | Adaptive Lasso |
| --- | --- | --- | --- | --- |
| $p$-value $< 0.001$ | 125 (42.8%) | 63 (21.6%) | 95 (32.5%) | 31 (10.6%) |
| $p$-value $< 0.005$ | 189 (64.7%) | 98 (33.6%) | 135 (46.2%) | 63 (21.6%) |
| $p$-value $< 0.01$ | 208 (71.2%) | 117 (40.1%) | 158 (54.1%) | 90 (30.8%) |
| $p$-value $< 0.05$ | 249 (85.3%) | 159 (54.5%) | 225 (77.1%) | 155 (53.1%) |

Ridge and adaptive lasso methods showed similar overall sensitivity at *p*-value *<* 0.05, identifying 159 (54.5%) and 155 (53.1%) significant pathways respectively. However, their performance differed substantially at stricter thresholds. The ridge method identified 63 (21.6%) pathways at *p*-value *<* 0.001, while the adaptive lasso detected 31 (10.6%) pathways at this threshold, indicating that adaptive lasso is more conservative for highly significant associations.

All methods exhibited an expected pattern of decreasing detection rates as the significance threshold became more stringent, but the rate of decrease varied among methods. The difference in detection rates between methods was most pronounced at stricter thresholds, particularly between shrinkage and adaptive lasso methods. For instance, at *p*-value *<* 0.005, shrinkage identified 189 (64.7%) pathways while adaptive lasso detected 63 (21.6%) pathways, highlighting substantial differences in sensitivity between these approaches.

These findings largely align with our simulation results, with the shrinkage estimator maintaining its superior performance in both settings. The simulation study showed the shrinkage method achieving robust performance across different scenarios, which was confirmed in the pathway application, where it identified the highest number of significant pathways. However, while graphical lasso showed concerning performance in simulations, particularly in unbalanced scenarios, it showed strong pathway detection ability in this balanced real data application. Ridge and adaptive lasso methods showed more conservative behavior, with notably lower detection rates at stringent thresholds, reflecting their simulation performance patterns. These results suggest that while simulation studies provide valuable insights into method performance under controlled conditions, the behavior of these methods in real applications may show some variations due to the inherent complexity of biological systems and the influence of data balance.

## 5 Clinical validation of LCT results

While GSA outputs lists of gene sets deemed as statistically significant for a given condition, validating them is often a challenge as the ground truth on their significance could finally be provided by biological experiments, In addition to our use of use of Jaccard Index as a baseline threshold of disease relevance of a gene set, here we validate the LCT results based on clinically observed SLE-relevant phenotypes listed in the Human Phenotype Ontology (HPO) database. HPO also curated a gene set associated with each phenotype. In our validation experiment, we selected a collection of 5 SLE-relevant phenotypes, and to compare, 5 matched phenotypes that are not known to be associated with SLE. These clinical phenotypic gene sets were used for running LCT on the SLE gene expression study GSE38351 involving 14 SLE cases and 12 healthy controls. The LCT results shown in Table 3 clearly demonstrate the high significance of the SLE-relevant phenotypes as opposed to the statistical insignificance (using LCT p-value threshold 0.05) of the SLE-irrelevant ones.

**Table 3.** Clinical validation of LCT results.

| Clinical Phenotype | HPO Id | SLE | LCT |
| --- | --- | --- | --- |
|  |  | Relevance | P-value |
| Discoid lupus rash | HP:0007417 | 1 | <0.001 |
| Scaling skin | HP:0040189 | 0 | 0.116 |
| Malar rash | HP:0025300 | 1 | <0.001 |
| Squamous cell carcinoma of the skin | HP:0006739 | 0 | 0.06 |
| Glomerulonephritis | HP:0000099 | 1 | 0.003 |
| Multiple glomerular cysts | HP:0100611 | 0 | 0.733 |
| Antinuclear antibody positivity | HP:0003493 | 1 | <0.001 |
| Anti-acetylcholine receptor antibody positivity | HP:0030208 | 0 | 0.089 |
| Oral ulcer | HP:0000155 | 1 | <0.001 |
| Oral leukoplakia | HP:0002745 | 0 | 0.348 |

*Note:* In the SLE Relevance column, 1 indicates a phenotype that is clinically relevant to SLE, and 0 indicates a phenotype not typically associated with SLE. HPO stands for Human Phenotype Ontology.

## 6 Discussion

The present work focused on two important aspects of GSA: (a) assessing the performance of various covariance matrices as used in LCT for testing the association of expression of genes from a set and a binary outcome, and (b) checking the robustness of LCT in the presence of unbalanced designs, which are not uncommon in biomedical studies, based on real datasets. We are not aware of any other simulation-based or comparative analysis for evaluating the performance of GSA methods for balanced versus unbalanced gene expression studies. While there are several GSA methods in the literature, here we chose LCT to investigate the performance of the covariance matrix options, as well as its robustness to the analysis of unbalanced datasets in comparison with GSEA.

In the past, LCT Type I error, power and computational efficiency were compared against top GSA methods in simulations and real data analysis studies (Wang et al., 2014). The results of our previously published simulation studies indicated that LCT Type I error and power were comparable to MANOVA-GSA (Tsai and Chen, 2009), and superior to SAM-GS (Dinu et al., 2007), especially at higher magnitudes of the correlation values across gene sets, which is a common scenario in GSA. Due to an algebraic property of the eigenvalue decomposition of the shrinkage covariance matrix estimator, LCT was superior to both methods in terms of computational efficiency (Wang et al., 2011). Moreover, LCT outperformed GSEA in a previous simulation study by Khodayari Moez et al. (2019), which the present study extended to gene expression data with unbalanced sample sizes.

Interestingly, Goeman et al. (2004) discouraged comparing GSA methods across two different categories that essentially test different hypotheses. These categories are known as (1) self-contained GSA methods – which include only genes in a pre-defined set and use a traditional approach to evaluate the significance of the association between the set and a binary phenotype (when the statistical test is not asymptotically tractable, the phenotypic group labels are permuted); and (2) competitive GSA methods – which involve genes outside the pre-defined set of genes, and use a permutation of the genes in the list to evaluate the association of the pre-defined set with the binary outcome, therefore assuming the gene expression measurements across a pre-defined set are independent. Goeman and Bühlmann (2007) pointed out in their methodological review that this independence assumption is not tenable since genes are gathered into pre-defined sets on the premise that they share common biological functions.

We understand that our study has certain limitations. While gene expression studies could use various experimental designs, we have focused here only on a very specific, albeit common, design issue. Further, we primarily focused on LCT as our chosen GSA method mainly because of its variety of past successful applications (as outlined in the Introduction). This could be addressed by a more general comparative study in the future. For our comparative analysis, we included the popular method GSEA for both the real data applications. The comparison of the performances of LCT and GSEA demonstrated that the former gives more powerful and robust results for these datasets, which were found to be consistent with our previous simulation studies (Dinu et al., 2007; Wang et al., 2014). In addition to identifying more gene sets that are relevant to the disease under investigation, LCT does so consistently for both larger as well as smaller sample sizes that characterized the two datasets, while GSEA seemed to have difficulty in doing this for the smaller sample size scenario corresponding to the second dataset.

Researchers should exercise particular caution when applying LCT to highly unbalanced study designs. Our findings reveal that when the case-to-control ratio exceeds 3:1, the default shrinkage estimator exhibits elevated Type I error rates, particularly with high correlation across the gene set. This highlights the need for more conservative significance thresholds. Additionally, ridge estimation, while more robust, may reduce the power to detect subtle gene set associations.

## 7 Acknowledgement

The authors declare that there are no conflicts of interest.

## References

Amini, P., Hajihosseini, M., Pyne, S., Dinu, I.: Geographically weighted linear combination test for gene-set analysis of a continuous spatial phenotype as applied to intratumor heterogeneity. Frontiers in Cell and Developmental Biology 11, 1065586 (2023)

Dinu, I., Moez, E.K., Hajihosseini, M., Leite, A.P., Pyne, S.: Use of linear combination test to identify gene signatures of human embryonic development in single cell rna-seq experiments. Statistics and Applications 19(1), 431–442 (2021)

Dinu, I., Potter, J.D., Mueller, T., Liu, Q., Adewale, A.J., Jhangri, G.S., Einecke, G., Famulski, K.S., Halloran, P., Yasui, Y.: Improving gene set analysis of microarray data by sam-gs. BMC Bioinformatics 8, 1–13 (2007)

Dinu, I., Wang, X., Kelemen, L.E., Vatanpour, S., Pyne, S.: Linear combination test for gene set analysis of a continuous phenotype. BMC Bioinformatics 14, 1–9 (2013)

Friedman, J., Hastie, T., Tibshirani, R.: Sparse inverse covariance estimation with the graphical lasso. Biostatistics 9(3), 432–441 (2008)

Friedman, J., Hastie, T., Tibshirani, R.: Glasso: Graphical Lasso: Estimation of Gaussian Graphical Models. (2019). R package version 1.11. https://CRAN.R-project.org/package=glasso

Goeman, J.J., Bühlmann, P.: Analyzing gene expression data in terms of gene sets: methodological issues. Bioinformatics 23(8), 980–987 (2007)

Goeman, J.J., Van De Geer, S.A., De Kort, F., Van Houwelingen, H.C.: A global test for groups of genes: testing association with a clinical outcome. Bioinformatics 20(1), 93–99 (2004)

Hung, T., Pratt, G., Sundararaman, B., Townsend, M., Chaivorapol, C., Bhangale, T., Graham, R., Ortmann, W., Criswell, L., Yeo, G., et al.: The ro60 autoantigen binds endogenous retroelements and regulates inflammatory gene expression. Science 350(6259), 455–459 (2015)

Human MSigDB Collections: KEGG LEGACY Subset of CP from Human MSigDB v2023.2.Hs Collections. https://www.gsea-msigdb.org/gsea/msigdb/collections.jsp, Last accessed on 2024-07-03 (2023)

Kerr, K.F.: Comments on the analysis of unbalanced microarray data. Bioinformatics 25(16), 2035–2041 (2009)

Kanehisa, M., Furumichi, M., Sato, Y., Ishiguro-Watanabe, M., Tanabe, M.: Kegg: integrating viruses and cellular organisms. Nucleic Acids Research 49(D1), 545–551 (2021)

Kanehisa, M., Goto, S.: Kegg: Kyoto encyclopedia of genes and genomes. Nucleic Acids Research 28(1), 27–30 (2000)

Köhler, S., Gargano, M., et al.: The human phenotype ontology in 2021. Nucleic Acids Research 49, 1207–1217 (2021)

Kanehisa, M., Goto, S., Sato, Y., Furumichi, M., Tanabe, M.: KEGG for integration and interpretation of large-scale molecular data sets. Accessed: 2024-06-30 (2012). https://www.kegg.jp/

Khodayari Moez, E., Hajihosseini, M., Andrews, J.L., Dinu, I.: Longitudinal linear combination test for gene set analysis. BMC Bioinformatics 20, 1–19 (2019)

Khodayari, E., Pyne, S., Dinu, I.: Association between bivariate expression of key oncogenes and metabolic phenotypes of patients with prostate cancer. Computers in Biology and Medicine 103, 55–63 (2018)

Lin, W.-J., Hsueh, H.-M., Chen, J.J.: Power and sample size estimation in microarray studies. BMC Bioinformatics 11, 1–9 (2010)

Mootha, V.K., Lindgren, C.M., Eriksson, K.-F., Subramanian, A., Sihag, S., Lehar, J., Puigserver, P., Carlsson, E., Ridderstråle, M., Laurila, E., et al.: Pgc-1α-responsive genes involved in oxidative phosphorylation are coordinately downregulated in human diabetes. Nature genetics 34(3), 267–273 (2003)

Opgen-Rhein, R., Strimmer, K.: Accurate ranking of differentially expressed genes by a distribution-free shrinkage approach. Statistical applications in genetics and molecular biology 6(1) (2007)

Peeters, C.F.W., Bilgrau, A.E., van Wieringen, W.N.: rags2ridges: A one-stop-l2-shop for graphical modeling of high-dimensional precision matrices. Journal of Statistical Software 102(4), 1–32 (2022) 10.18637/jss.v102.i04

Rothman, A.J., Levina, E., Zhu, J.: Generalized thresholding of large covariance matrices. Journal of the American Statistical Association 104(485), 177–186 (2009)

RStudio team: RStudio: Integrated Development Environment for R. RStudio, PBC., Boston, MA (2023). RStudio, PBC. http://www.rstudio.com/

Smiljanovic, B., Grün, J.R., Biesen, R., Schulte-Wrede, U., Baumgrass, R., Stuhlmüller, B., Maslinski, W., Hiepe, F., Burmester, G.-R., Radbruch, A., et al.: The multifaceted balance of tnf-α and type i/ii interferon responses in sle and ra: how monocytes manage the impact of cytokines. Journal of molecular medicine 90, 1295–1309 (2012)

Schafer, J., Opgen-Rhein, R., Zuber, V., Ahdesmaki, M., Silva, A.P.D., Strimmer., K.: Corpcor: Efficient Estimation of Covariance and (Partial) Correlation. (2021). R package version 1.6.10. https://CRAN.R-project.org/package=corpcor

Schäfer, J., Strimmer, K.: A shrinkage approach to large-scale covariance matrix estimation and implications for functional genomics. Statistical Applications in Genetics and Molecular Biology 4(1) (2005)

Subramanian, A., Tamayo, P., Mootha, V.K., Mukherjee, S., Ebert, B.L., Gillette, M.A., Paulovich, A., Pomeroy, S.L., Golub, T.R., Lander, E.S., et al.: Gene set enrichment analysis: a knowledge-based approach for interpreting genome-wide expression profiles. Proceedings of the National Academy of Sciences 102(43), 15545–15550 (2005)

Tsai, C.-A., Chen, J.J.: Multivariate analysis of variance test for gene set analysis. Bioinformatics 25(7), 897–903 (2009)

Van Wieringen, W.N., Peeters, C.F.: Ridge estimation of inverse covariance matrices from high-dimensional data. Computational Statistics & Data Analysis 103, 284–303 (2016)

Wang, X., Dinu, I., Liu, W., Yasui, Y.: Linear combination test for hierarchical gene set analysis. Statistical Applications in Genetics and Molecular Biology 10(1) (2011)

Wang, X., Pyne, S., Dinu, I.: Gene set enrichment analysis for multiple continuous phenotypes. BMC Bioinformatics 15, 1–9 (2014)

Wither, J.E., Prokopec, S.D., Noamani, B., Chang, N.-H., Bonilla, D., Touma, Z., Avila-Casado, C., Reich, H.N., Scholey, J., Fortin, P.R., et al.: Identification of a neutrophil-related gene expression signature that is enriched in adult systemic lupus erythematosus patients with active nephritis: Clinical/pathologic associations and etiologic mechanisms. PloS One 13(5), 0196117 (2018)

Yang, K., Li, J., Gao, H.: The impact of sample imbalance on identifying differentially expressed genes. BMC Bioinformatics 7, 1–13 (2006)

Zou, H.: The adaptive lasso and its oracle properties. Journal of the American statistical association 101(476), 1418–1429 (2006)

